# Identification of NAD-RNAs and ADPR-RNA decapping in the archaeal model organisms *Sulfolobus acidocaldarius* and *Haloferax volcanii*

**DOI:** 10.1101/2022.11.02.514978

**Authors:** José Vicente Gomes-Filho, Ruth Breuer, Hector Gabriel Morales-Filloy, Nadiia Pozhydaieva, Andreas Borst, Nicole Paczia, Jörg Soppa, Katharina Höfer, Andres Jäschke, Lennart Randau

**Affiliations:** Faculty of Biology, Philipps-Universität Marburg, Marburg, Germany; Institute of Pharmacy and Molecular Biotechnology (IPMB), Heidelberg University, Heidelberg, Germany; Institute for Molecular Biosciences, Biocentre, Goethe-University, Frankfurt, Germany; Max Planck Institute for Terrestrial Microbiology, Marburg, Germany; SYNMIKRO, Center for Synthetic Microbiology, Marburg, Germany

**Keywords:** Archaea, NAD, ADP-ribose, 5’-RNA caps, Transcriptome, RNA modification, hyperthermophiles

## Abstract

NAD is a coenzyme central to metabolism that was also found to serve as a 5’-terminal cap of bacterial and eukaryotic RNA species. The presence and functionality of NAD-capped RNAs (NAD-RNAs) in the archaeal domain remain to be characterized in detail. Here, by combining LC-MS and NAD captureSeq methodology, we quantified the total levels of NAD-RNAs and determined the identity of NAD-RNAs in the two model archaea, *Sulfolobus acidocaldarius* and *Haloferax volcanii*. A complementary differential RNA-Seq (dRNA-Seq) analysis revealed that NAD transcription start sites (NAD-TSS) correlate with well-defined promoter regions and often overlap with primary transcription start sites (pTSS). The population of NAD-RNAs in the two archaeal organisms shows clear differences, with *S. acidocaldarius* possessing more capped small non-coding RNAs (sncRNAs) and leader sequences. The NAD-cap did not prevent 5’→3’ exonucleolytic activity by the RNase Saci-aCPSF2. To investigate enzymes that facilitate the removal of the NAD-cap, four Nudix proteins of *S. acidocaldarius* were screened. None of the recombinant proteins showed NAD decapping activity. Instead, the Nudix protein Saci_NudT5 showed activity after incubating NAD-RNAs at elevated temperatures. Hyperthermophilic environments promote the thermal degradation of NAD into the toxic product ADPR. Incorporating NAD into RNAs and the regulation of ADPR-RNA decapping by Saci_NudT5 is proposed to provide additional layers of maintaining stable NAD levels in archaeal cells.

**Importance:** This study reports the first characterization of 5’-terminally modified RNA molecules in Archaea and establishes that NAD-RNA modifications, previously only identified in the other two domains of life, are also prevalent in the archaeal model organisms *Sulfolobus acidocaldarius* and *Haloferax volcanii*. We screened for NUDIX hydrolases that could remove the NAD-RNA cap and showed that none of these enzymes removed NAD modifications, but we discovered an enzyme that hydrolyzes ADPR-RNA. We propose that these activities influence the stabilization of NAD and its thermal degradation to potentially toxic ADPR products at elevated growth temperatures.

## Introduction

The discovery of NAD, a cofactor critical to cellular metabolism, as a 5’ cap in Bacteria challenged earlier notions that only Eukaryotes utilize RNA capping mechanisms (1). Since the first discovery of NAD-RNA caps, additional reports of their presence in Gram-positive bacteria and Eukaryotes such as *Arabidopsis thaliana, Saccharomyces cerevisiae*, and mammalian cells suggest that this RNA modification is ubiquitous in the tree of life (2–8). The reported concentrations of NAD covalently linked to RNAs are variable and range from 1.9 fmol / μg in human cells to 116 fmol / μg in the stationary growth phase of *Mycobacterium smegmatis* (9). Mechanistic studies demonstrated that bacterial RNA polymerase (RNAP) and eukaryotic RNAP II can utilize NAD, NADH, Flavin Adenine Dinucleotide (FAD), Adenosine diphosphate ribose (ADPR), and 3’-dephospho-coenzyme A (dpCoA) to initiate transcription at promoters containing A at its +1 position (4, 10, 11). Additionally, the presence of NAD-caps on mammalian small nucleolar RNAs (snoRNAs) and the related small Cajal body RNAs (scaRNAs) suggests the presence of an additional post-transcriptional capping mechanism in eukaryotic cells (6).

In human and fungal cells, the non-canonical decapping enzymes DXO/Rat1 are responsible for initiating NAD-RNA degradation by removing the NAD-cap (6, 3). In *Escherichia coli*, NAD decapping is performed by a nucleoside diphosphate linked to another moiety X (NUDIX) protein termed NudC (12). This enzyme hydrolyses the NAD-cap resulting in nicotinamide mononucleotide (NMN) and RNA with a 5’-monophosphate terminus (5’-p-RNA) that is efficiently degraded by cellular 5’→3’ exonucleases (12, 8, 13). Further studies aiming to elucidate the function of NAD-RNAs revealed striking differences in the roles of this modification in bacterial and eukaryotic cells. In *E*.*coli*, it was initially thought that this modification could protect the

RNA against pyrophosphohydrolase (RppH) and RNAse E degradation, but more recent *in vitro* studies argue that RppH also functions as a NAD decapping enzyme (13). In *Bacillus subtillis*, it was shown that NAD modification of RNAs prevents 5’→3’ exonucleolytic activity from RNase J1, suggesting a stabilizing role (8). On the other hand, in eukaryotic cells, NAD-caps are considered to promote RNA decay (6), and a highly efficient surveillance machinery for the degradation of NAD-RNAs was described for yeast (14). The presence of NAD-caps can be related to different biological outcomes, even in organisms from the same domain of life, as demonstrated by the putative translational capacity of NAD-RNAs in eukaryotic cells (6, 5). Moreover, the 5’→3’ exonucleases Xrn1 and Rat1 from yeast mitochondria are suggested to directly influence the concentration of free NAD by releasing intact NAD from NAD-RNAs (14).

The degradation of NAD at high temperatures (>75°C) into nicotinamide (Nm) and ADPR demands hyperthermophilic microorganisms to present robust pathways for detoxifying these products (15, 16). In mesophilic organisms, the generation of ADPR is mainly achieved through enzymatic reactions performed by enzymes such as ADPR-transferases, cyclic ADPR-synthases, and poly ADPR polymerases (17). A recent study provided the first evidence for *in vivo* 5’ ADP-ribosylated RNAs (ADPR-RNA) in mammalian cells (18). Interestingly, the process of ADPR-capping in Eukaryotes has different pathways. The human protein CD38, for example, can convert NAD-RNA to ADPR-RNA by removing an Nm from the NAD-generating RppAp-RNA (19). The bacterial RNA 2’-phosphotransferase (Tpt1) and its orthologues from higher organisms, TRPT1, can use free NAD to ADP-ribosylate 5’-p-RNA substrates and, unlike CD38, this process generates ApRpp-RNA (20, 18). In both cases, contrary to NAD-capping, the generation of ADPR-RNAs is achieved post-transcriptionally. Furthermore, ADPR-RNAs were shown to be more resistant to Xrn1 exonuclease activity while not supporting translation (18).

As we gain first insights into the functional consequences of NAD-capping in Bacteria and Eukaryotes, this information is lacking for the Archaea. Here, we combine LC-MS and NAD captureSeq methodologies to quantify NAD-RNA levels and determine the identity of NAD-capped RNAs in the two archaeal model organisms, *Sulfolobus acidocaldarius* and *Haloferax volcanii*. Multiple NUDIX family proteins can be involved in processing mRNA caps (21). A sequence similarity search provided four NUDIX protein candidates in *S. acidocaldarius. In vitro* assays using recombinant enzymes did not reveal NAD decapping activity. Instead, we detected that SACI_RS00060 (here renamed to Saci_NudT5) showed activity following heat exposure of NAD-capped RNAs. We propose that thermal degradation generates RppAp-RNA (now referred to as ADPR-RNA) substrates for this enzyme and suggest that NAD-capping influences the thermal stabilization of NAD in *S. acidocaldarius* and other hyperthermophilic organisms.

## Results

### Detection and quantification of NAD-capped RNAs in *S. acidocaldarius* and *H. volcanii*

First, we aimed to determine NAD modifications of RNAs in the crenarchaeon *S. acidocaldarius* and the euryarcheon *H. volcanii*. Nuclease P1 is a 3’→5’ exonuclease that releases single nucleotides without affecting pyrophosphate bonds, leaving capping nucleotides, like NAD, intact after release (Fig. 1A). Total RNA was isolated from *S. acidocaldarius* and *H. volcanii* and treated with nuclease P1. NAD was identified by detecting the compound-specific mass transitions 662 (m/z) → 540 (m/z) and 662 (m/z) → 273 (m/z) (Fig. 1B and C). To determine the levels of co-purified free NAD, we analyzed RNA treated with heat-inactivated nuclease P1 (Second peak profile in Fig. 1B and 1C). In addition, standards ranging from 0.1 nM to 1 μM were used to calculate the total concentration of NAD released after nuclease P1 digestion (Fig. 1B and C, third panel). After normalizing to RNA mass (100 μg), the determined concentrations of NAD were 260±72 fmol per μg RNA for *S. acidocaldarius* and 110±9 fmol per μg RNA for *H. volcanii*. These results indicate the presence of NAD-RNAs in Archaea and establish *S. acidocaldarius* as the organism with the highest concentration of NAD-RNAs detected so far.

**Figure 1:**
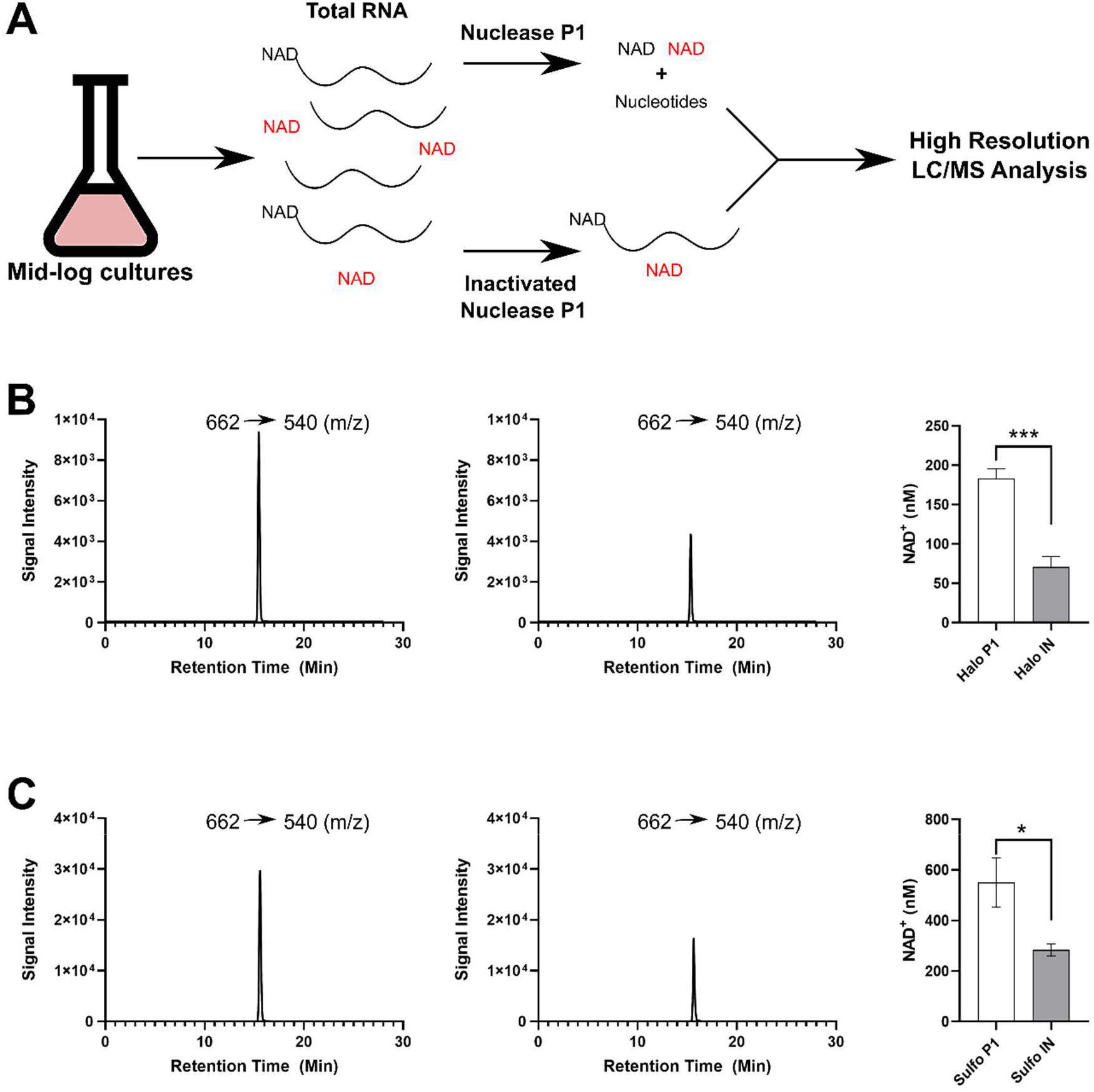
Detection of NAD in total RNA extract from *S. acidocaldarius* and *H. volcanii*. **A)** Method utilized to quantify free (red) or RNA-bound NAD (black). Briefly, total RNA is extracted from mid-log cultures and digested with either nuclease P1 or with a heat-inactivated enzyme. Next, samples are submitted to LC-MS analysis, and NAD is measured. **B)** Peak intensity of the NAD specific mass transition 662 (m/z) → 540 (m/z) for *H. volcanii* total RNA digested with nuclease P1, inactivated nuclease P1, and quantification results (P1: Treated RNA; IN: Inactivated nuclease P1 samples). **C)** Peak intensity of the NAD specific mass transition 662 (m/z) → 540 (m/z) for *S. acidocaldarius* total RNA digested with nuclease P1, Inactivated nuclease P1, and quantification results (P1: Treated RNA; IN: Inactivated nuclease P1 samples). The final concentration per μg of RNA was established after subtracting the background NAD from the RNA samples (Inactivated nuclease P1) and calculated using a calibration curve (0.1 nM to 1 μM NAD). Asterisks correspond to the t-test p-value (*: <0.05; ***: <0.01), n = 3.

### Identification and classification of NAD-RNAs in *S. acidocaldarius* and *H. volcanii*

To obtain a snapshot of the NAD-RNA populations from both organisms, total RNA was extracted during the mid-log growth phase, and NAD captureSeq libraries were prepared (1, 13). The libraries were sequenced on an Illumina HiSeq3000, and at least 6 million reads were obtained per sample. The obtained reads were trimmed and aligned against the archaeal genomes of interest. Next, DESeq2 (22) was used to determine enriched transcripts (p-adjusted value < 0.1 and log2(Fold Change) > 1) in samples treated with an ADP-ribosyl cyclase from *Aplysia california* (ADPRC+) versus non-treated samples (ADPRC-) (13, 1). Using these threshold values, we identified 86 NAD-RNAs for *H. volcanii* and 83 NAD-RNAs for *S. acidocaldarius* (Supplementary Tables S1 and S2). We used previously published data to compare the 50 most abundant transcripts in the ADPRC+ libraries with the 50 most abundant transcripts in an sRNA-seq library obtained under identical growth conditions (23). From the enriched RNAs, only 6 were amongst the most expressed in *S. acidocaldarius* (Supplementary Table 3 and Supplementary Figure 1). To further confirm the calculated enrichment in our datasets, qPCR analysis was performed with cDNA obtained after ligation of the second adapter. This experiment had two enriched genes, *tfb* and SACI_RS10480, and one negative control, SACI_RS00345, as targets. In agreement with the enrichment detected by NAD captureSeq, *tfb* and SACI_RS10480 showed a relative expression of 25±10 and 45±20, respectively. The negative control SACI_RS00345 did not show any enrichment. Thus, the sRNA-seq analysis and the qPCR validation reinforce a selective enrichment of NAD-RNAs and not a bias for overly abundant transcripts. Analysis of nucleotide frequency of the +1 NAD transcription start sites (NAD-TSS) and -1 positions demonstrated that all enriched transcripts start with an adenine (Fig. 2A and B). For both *H. volcanii* and *S. acidocaldarius*, the -1 position was found to be enriched for thymine. For positions -2 to -3, *S. acidocaldarius* presented a slight preference for A/T compared to G/A in *H. volcanii* (Fig. 2A and B). To further evaluate if the addition of the NAD-cap occurs co-transcriptionally, we analyzed the upstream regions (-50 bp) of the identified NAD-RNAs for recognizable promoter motifs. A TFB recognition element (BRE) was detected at around position -30 for *S. acidocaldarius*. A TATA box motif was also detected at around position -26 for *S. acidocaldarius* and position -28 for *H. volcanii* (Fig. 2A and B) (24, 25). Next, we sought to compare the NAD-TSS with the primary transcription start sites (pTSS) containing a 5’-ppp. To this end, we prepared dRNA-Seq libraries for *S. acidocaldarius* (Supplementary Table 4) and reanalyzed previously published data for *H. volcanii* (26). Manual curation of the positions showed that most NAD-TSS and pTSS are found at the same positions (76% for *S. acidocaldarius* and 90% for *H. volcanii*) (Fig. 2C). The high number of overlapping NAD-TSSs and pTSSs, together with the detection of distinct promoter motifs, further supports that archaeal RNAs are co-transcriptionally capped with NAD (10, 27).

**Figure 2:**
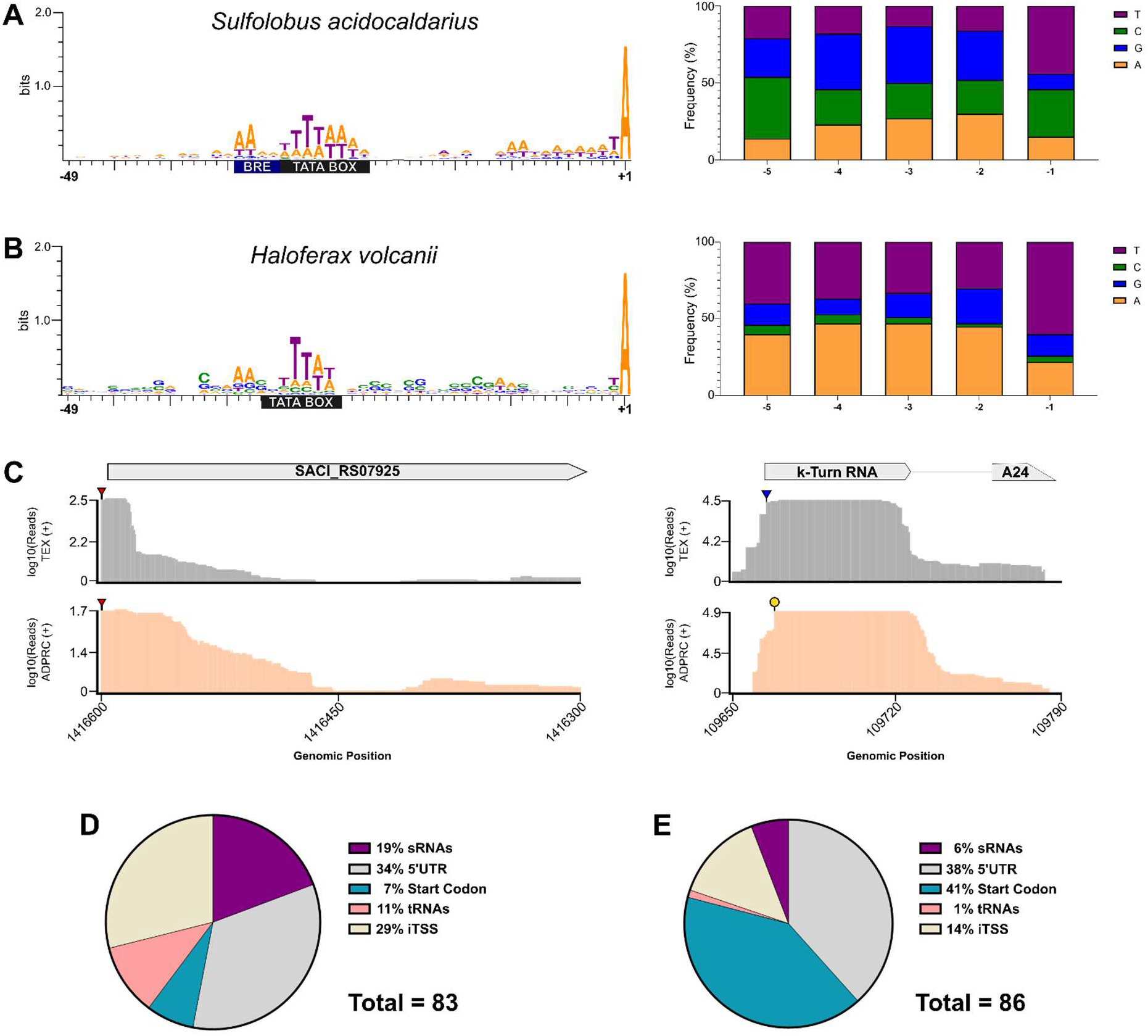
Promoter identification, nucleotide frequency of the -1 to -5 positions relative to NAD-RNAs, and primary transcription start sites (pTSS) comparison. A) Promoter and nucleotide frequency analysis for *S. acidocaldarius*. The blue rectangle represents the TFB recognition element (BRE). B) Promoter and nucleotide frequency analysis for *H. volcanii*. C) Comparison of transcription starts sites identified by dRNA-Seq (grey lines) and NAD captureSeq (salmon lines). Left panel: Coverage plot of carboxypeptidase M32 (SACI_RS07925) with matching NAD- and pTSS (red triangles). Right panel: Coverage plot of a k-turn RNA upstream of the peptidase A24 with non-matching NAD-TSS (yellow circle) and pTSS (blue triangle). Classification of NAD-RNAs identified in *S. acidocaldarius* (D) and in *H. volcanii* (E).

To explore potential patterns of functional enrichment of capped RNA molecules, the identified NAD-RNAs were divided into five categories: I) Internal: the +1 position is located within a coding gene; II) Start codon: the +1 position matches the annotated start codon; III) tRNAs; IV) small RNAs (sRNAs, e.g., C/D box sRNAs and non-coding sRNAs); V) 5’ UTRs: the +1 position is located upstream of the start codon of an enriched coding gene (Fig. 2D and E). Comparing the abundances of each class between *H. volcanii* and *S. acidocaldarius* revealed some striking differences. First, 9 tRNAs were enriched in *S. acidocaldarius*, while only tRNA-Met was enriched in *H. volcanii* (Supplementary Tables 1 and 2). Second, only 7% of the enriched mRNAs for *S. acidocaldarius* were identified to contain NAD matching the start codon adenosine of the respective genes, as opposed to 41% in *H. volcanii*. The number of NAD-capped sRNAs detected in *S. acidocaldarius* was almost four times higher than for *H. volcanii*. Next, to obtain an overview of the enriched gene functions, Archaeal Clusters of Orthologous Genes (arCOGs) were used to group genes according to different biological functions (15). However, no clear enrichment of specific arCOGs was detected (data not shown).

In both archaea, the Transcription Initiation Factor IIB (TFIIB) enrichment was visualized, arguing for a conserved role of the NAD-cap in this gene’s transcript. Moreover, in *H. volcanii*, the mRNA of a NAD-dependent protein deacetylase gene from the SIR2 family can be NAD-capped, suggesting a possible connection between the intracellular levels of free NAD and NAD capping. Previous studies demonstrated that C/D box sRNAs are abundant in archaeal cells and are crucial for the guided methylation of RNA targets in *S. acidocaldarius* (23, 28). Thus, the NAD-cap might directly influence the stability of this subset of C/D box RNAs in *S. acidocaldarius*. In eukaryotic cells, the biogenesis of NAD-capped tRNAs and snoRNAs is still a point of contention. A previous study demonstrated that some NAD-capped snoRNAs and tRNAs did not possess a recognizable upstream sequence motif that supports NAD-initiated transcription (29). These studies raised the hypothesis that these candidates may be post-transcriptionally NAD-capped. In our *S. acidocaldarius* dataset, the analysis of the upstream region of NAD-capped tRNAs and snoRNAs evidenced the presence of recognizable promoter motifs, reinforcing that NAD-capping can also occur co-transcriptionally for these transcripts.

### The NAD-cap does not prevent Saci-aCPSF2 5’→3’exonucleolytic activity

The model archaeon *S. acidocaldarius* was used to investigate proteins that might influence NAD-RNA turnover. RNase J is a widespread exo/endoribonuclease in Bacteria and Archaea (30). In *B. subtillis*, the 5’→3’-exonucleolytic activity of RNase J1 relies on the presence of a monophosphate group at the 5’ end of different transcripts (8). Additionally, the NAD-cap was not as efficient as a 5’-ppp against RNase J1 activity (8). In *S. acidocaldarius*, the RNase J orthologue Saci-aCPSF2 was shown to act as an exonuclease against 5’-p-RNAs substrates while retaining some activity against 5’-ppp-RNAs (31, 32). Therefore, *in vitro* assays were performed to evaluate the impact of the NAD-cap on the exonucleolytic activity of recombinant Saci-aCPSF2. NAD-RNAs were not found to be protected against Saci-aCPSF2 activity but were instead a preferential substrate for degradation (Supplementary Figure. 4A and B).

### NUDIX proteins from *S. acidocaldarius* have ADPR-decapping activity but cannot perform NAD decapping

In Bacteria, the first identified NAD decapping enzyme was NudC, a member of the NUDIX family, which hydrolyses the NAD-cap resulting in 5’-p-RNA and free nicotinamide mononucleotide (NMN) (12). The family of Nudix hydrolases encompasses functionally diverse and versatile proteins, all containing the conserved Nudix motif with the consensus sequence GX_5_EX_5_U/AXREX_2_EEXGU (U for hydrophobic residue and X for any residue) (21). More recently, both *E. coli* RppH and *Bacillus subtilis* BsRppH were also shown to perform *in vitro* NAD decapping in addition to their pyrophosphohydrolase activities (8). Using a diverse set of Nudix proteins as template sequences to search for potential homologs in *S. acidocaldarius* yielded 4 protein candidates (Fig. 4A, Supplementary File 1). All four candidate proteins (SACI_RS00060, SACI_RS00575, SACI_RS00730, and SACI_RS02625) possess the conserved glutamic acid residues in the Nudix motif, which are crucial to the hydrolase activity (33, 12, 34, 21, 35). Another notable feature is the residue at position 16 following the G of the Nudix motif. The residue at this position was shown to suggest a possible substrate for the respective Nudix protein and therefore serves to identify and distinguish different subsets of Nudix hydrolases. In SACI_RS00060, a proline at this position suggests ADPR hydrolysis activity, while in SACI_RS00575, the tyrosine hints at activity towards polyphosphate dinucleoside substrates (35). For SACI_RS00730 and SACI_RS02625, no residue pointing at a specific activity was identified at this position.

Next, combining heterologous expression in *E. coli* and *in vitro* cell-free protein synthesis, we produced and purified these 4 identified Nudix proteins and generated individual Nudix domain mutants (NDM) (Supplementary Fig. 5). To evaluate the NAD decapping activity of these proteins, a synthetic RNA (model-RNA), containing a single A at its transcription start site, was *in vitro* transcribed using NAD, GTP, CTP, and UTP. The substitution of ATP for NAD ensures that the *in vitro* transcription reaction only initiates with the latter, providing pure NAD-RNA substrates. It was found that none of the recombinant *S. acidocaldarius* Nudix proteins performed NAD decapping *in vitro* (Fig. 3C), suggesting that either *S. acidocaldarius* has no enzymatic NAD decapping activity or another pathway is responsible for this process. As NAD was shown to be converted into ADPR and Nm at higher temperatures (see below), we continued to investigate if any *S. acidocaldarius* recombinant Nudix proteins could hydrolyze ADPR-RNA instead of NAD-RNAs. It was previously demonstrated that the Human NudT5 (HNudT5) hydrolyzes free ADPR *in vitro* (34). Additionally, through sequence analysis, SACI_RS00060 (here renamed to Saci_NudT5) clusters with other known ADPR-hydrolases, including HNudT5 (Fig. 3B). This led us to test the activity of these proteins against ADPR-RNAs. To this end, pure ADPR-RNA substrates were generated as previously described for NAD-RNAs by exchanging NAD for ADPR in the *in vitro* transcription reaction. The application of ADPR-decapping assays revealed that Saci_NudT5 could convert ADPR-RNAs to 5’-p-RNAs (Fig. 3D). As *S. acidocaldarius* might lack proteins with known NADase activity, such as the human CD38 and or the TIR domain proteins from Bacteria (19, 36) an alternative pathway is likely involved in the *in vivo* formation of ADPR-RNAs.

**Figure 3:**
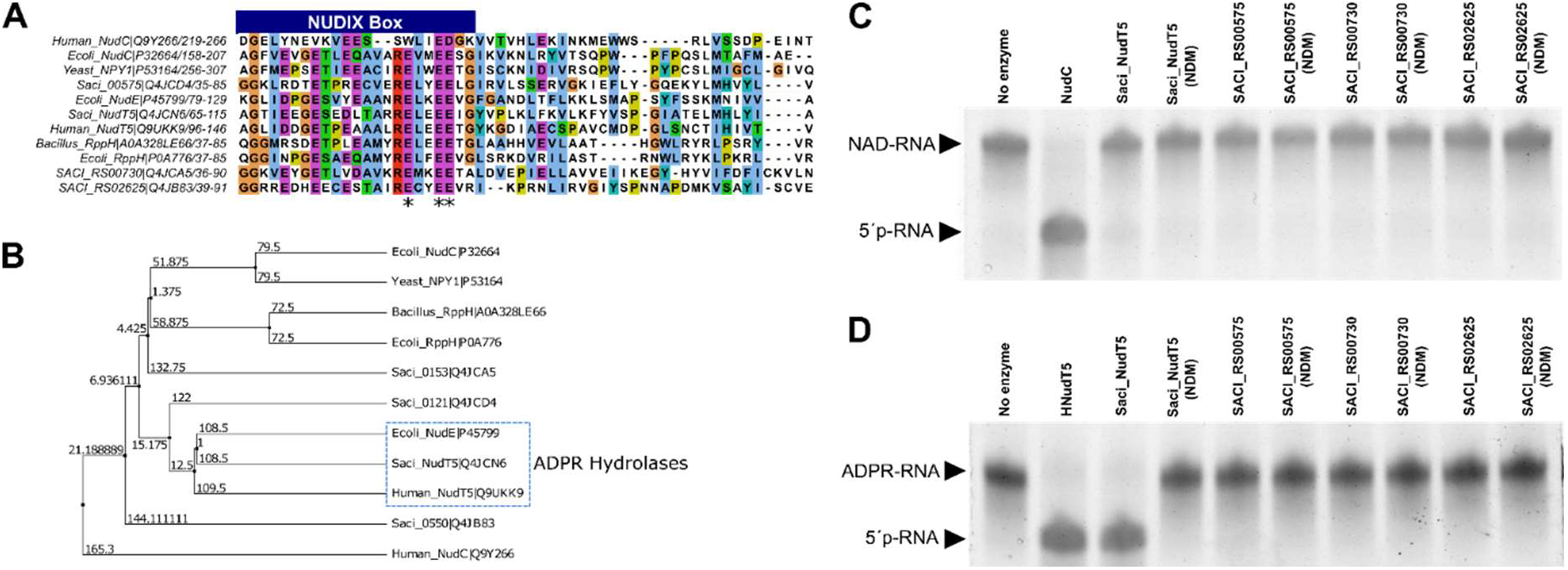
Identifying NUDIX proteins in *S. acidocaldarius* and evaluating NAD and ADPR-decapping activity. A) Alignment of NUDIX proteins of *S. acidocaldarius* and other organisms (Supplementary File 1). Blue rectangle: NUDIX Box motif. Asterisks: Selected amino acids to obtain NUDIX domain mutants (NDM) for each protein. B) Average distance tree using BLOSUM62 showing the grouping of Saci_NudT5 with the previously described ADPR-hydrolases NudE and HNudT5 (19, 51). C) NAD decapping activity of the four NUDIX candidates and their respective NDM was evaluated *in vitro* and resolved on APB-gels. D) ADPR-decapping activity of the four NUDIX candidates and their respective NDM was evaluated *in vitro* and resolved on APB-gels. Saci_NudT5 performed ADPR-decapping and Saci_NudT5 (NDM) lost this activity.

### NAD-RNAs are converted to ADPR-RNAs by thermal degradation

In thermophilic environments (> 60°C), such as the natural habitats of *S. acidocaldarius*, NAD is quickly degraded into ADPR and nicotinamide (Nm) (15, 37). Besides, hyperthermophilic Archaea contain robust pathways to salvage NAD from its degradation products (16, 15). A previous study demonstrated that the half-life of free NAD in 50 mM Tris-HCl buffer (pH 6.5 at 85°C) was 24 minutes at 85°C (15). We performed thermal degradation experiments to interrogate the stability of NAD covalently linked to RNAs (15). Briefly, *in vitro* transcribed model NAD-RNA was incubated at 75°C or 85°C in the presence of 50 mM Tris-HCl buffer (pH 6.5 at 85°C) for up to 2 hours. To track the conversion of NAD-RNA into ADPR-RNA, the heat-treated NAD-RNA was used for an ADPR-decapping assay with HNudT5, which shows *in vitro* activity toward ADPR but not NAD-RNAs (Fig. 4A and C) (34, 19). Interestingly, the obtained half-lives are significantly longer (54 and 50 minutes, for 75°C and 85°C, respectively) than what was previously determined for free NAD (Fig. 4B and D). Altogether, the high concentration of NAD-RNAs, the apparent absence of a NAD decapping enzyme, and the increased thermal stability of NAD covalently linked to RNA support that NAD-capping in *S. acidocaldarius* and possibly other hyperthermophilic organisms could have evolved to stabilize and store NAD, therefore slowing down the generation and accumulation of toxic compounds such as ADPR.

**Figure 4:**
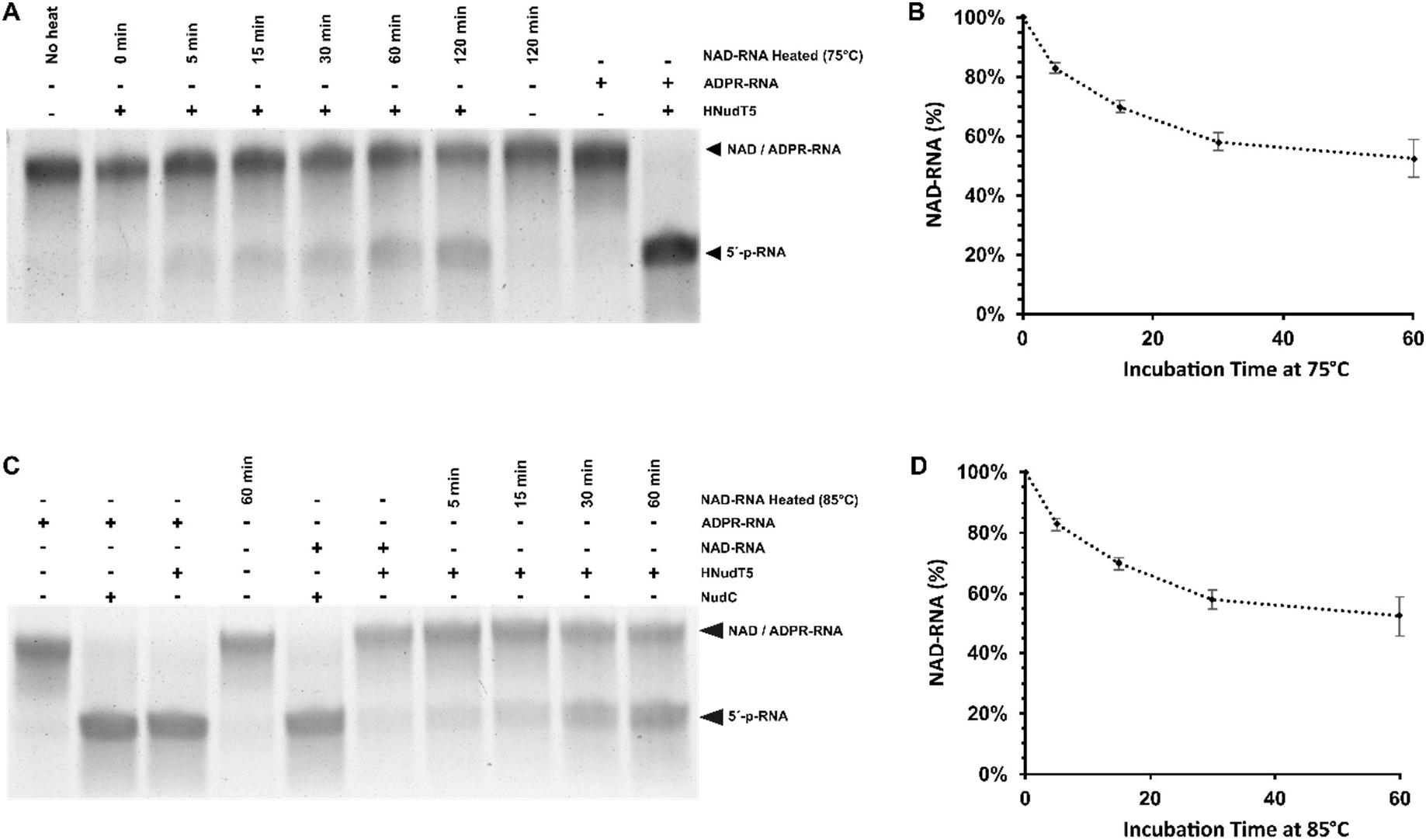
NAD-RNAs are converted to ADPR-RNAs by thermal degradation. A) NAD-RNAs were incubated at 75°C for up to 120 minutes in 50 mM Tris-HCl (pH 6.5 at 75°C). The reaction products were then incubated with HNudT5, and the conversion to 5’-p-RNA was monitored with APB-gels. *In vitro* transcribed NAD-RNA and ADPR-RNA were used as controls for HNudT5 reactions. B) Band intensities were used to calculate the ratio of NAD to ADPR-RNA conversion after heat treatment. C) NAD-RNAs were incubated at 85°C for up to 60 minutes in 50 mM Tris-HCl (pH 6.5 at 85°C). The reaction products were then incubated with HNudT5, and the conversion to 5’-p-RNA was monitored with APB gels. *In vitro* transcribed NAD-RNA and ADPR-RNA were used as controls for HNudT5 and NudC reactions. D) Band intensities were used to calculate the ratio of NAD to ADPR-RNA conversion after heat treatment. The t_1/2_ of NAD covalently linked to RNA was obtained with a typical decay equation (*dC/dt* = −*kdC*) (Mean ± SD, n = 3).

## Discussion

NAD and related dinucleotide metabolites are essential for many physiological processes, and their detection as 5’ caps for different bacterial and eukaryotic RNAs revealed additional layers of complexity (38). Nevertheless, the specific roles of NAD-caps are still being uncovered.

In the present study, the detection, quantification, and characterization of NAD-RNAs in the crenarchaeon *S. acidocaldarius* and in the euryarchaeon *H. volcanii* provided evidence that NAD-capping of RNA molecules is common to all domains of life.

None of the recombinant Nudix proteins from *S. acidocaldarius* exhibited NAD decapping activity, suggesting that this organism might utilize different pathways to process NAD-RNAs. Previous reports demonstrated that the non-canonical decapping enzymes DXO/Rai1 release intact NAD molecules from NAD-RNAs (6, 3, 14). Additionally, the highly conserved 5’-monophosphate 5’→3’ exoribonucleases, Xrn1 and Rat1, together with their interacting partner Rai1, can associate and hydrolyze NAD-RNAs *in vitro* (14). A previous study in *B. subtillis* showed that the 5’→3’-exonucleolytic activity from RNase J1 can be reduced by the presence of a NAD-cap (8). In *S. acidocaldarius*, the RNase J orthologue Saci-aCPSF2 is a known exonuclease that digests 5’-p-RNAs while retaining some activity against 5’-ppp-RNAs (31, 32). Surprisingly, we found that the NAD-cap not only does not prevent the exonucleolytic activity but is instead a preferential substrate of Saci-aCPSF2, highlighting another difference between bacterial and archaeal NAD-RNA turnover. Thus, additional mechanistic studies are required to elucidate if Saci-aCPSF2 could release intact NAD molecules, as described for DXO/Rai1 homologs.

Our results revealed that *S. acidocaldarius* has the highest concentration of NAD covalently linked to RNA identified. It is worth noting that, in hyperthermophilic environments, such as the natural habitats of *S. acidocaldarius*, NAD is quickly degraded into ADPR and Nm (15, 16). Therefore, hyperthermophilic organisms must present robust mechanisms to prevent the accumulation of toxic compounds generated via thermal degradation of NAD. Here, we demonstrate that the half-life of NAD bound to RNA is 67± minutes at 75°C and 64±12 minutes at 85°C, evidencing its higher stability when compared to free NAD (24 minutes at 85°C) (15, 37). Furthermore, the identification of Saci_NudT5 as an ADPR-decapping enzyme suggests a scenario where NAD-RNAs are spontaneously converted to ADPR-RNAs by thermal degradation and further processed to 5’-p-RNAs by Saci_NudT5. Therefore, we propose that NAD-capping, ADPR-capping, and NAD metabolism are interconnected in *S. acidocaldarius* (Fig. 5).

**Figure 5:**
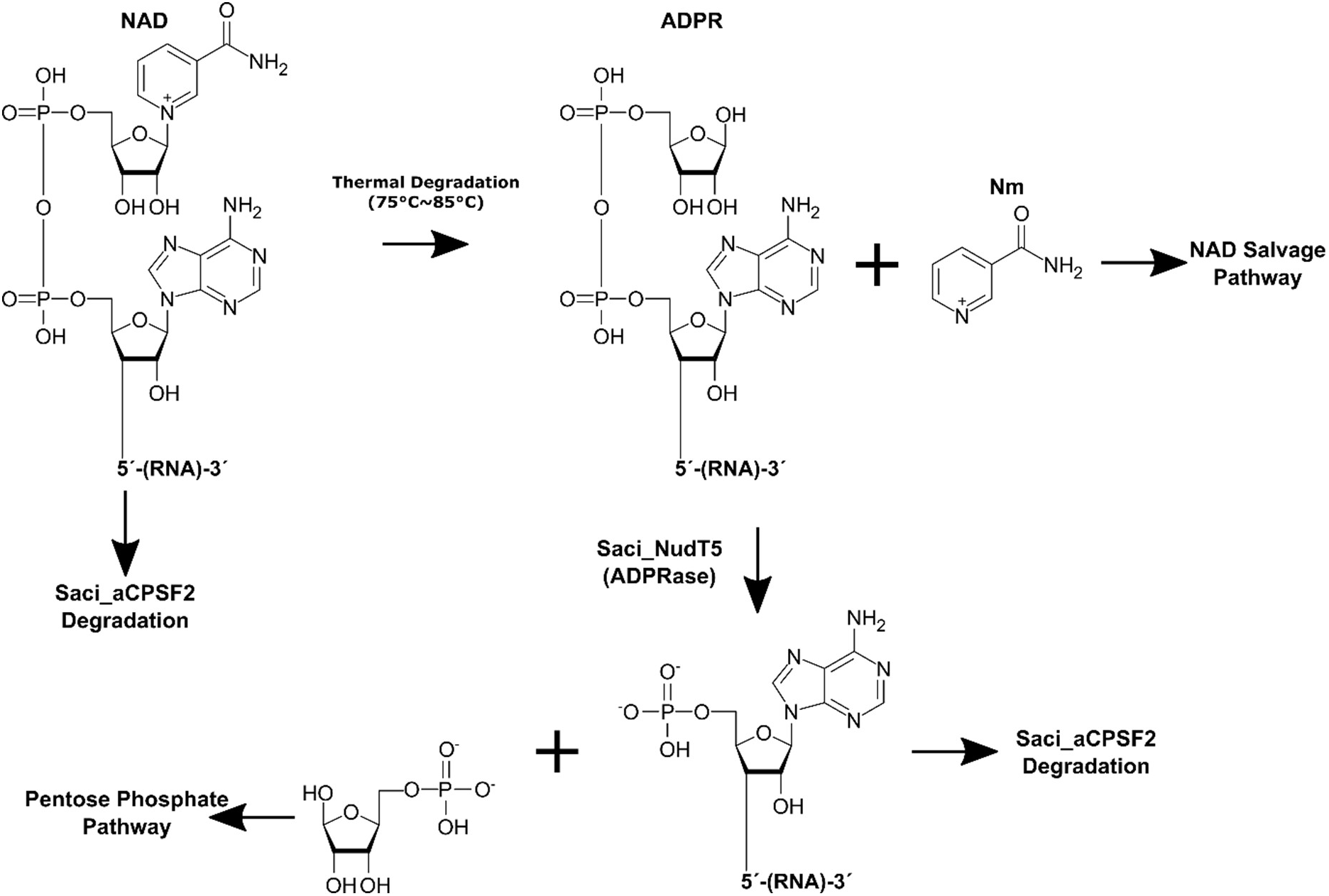
Proposed model for the relationship between NAD metabolism and RNA turnover *in S. acidocaldarius*. The thermal degradation of free NAD yields ADP-ribose and Nm. In *S. acidocaldarius*, the NAD salvage pathway is proposed to recover NAD from Nm. ADPR is converted to Ribose-5-Phosphate (R5P) and AMP by Saci_NudT5. These products can be utilized by the pentose phosphate pathway or for ATP synthesis. NAD molecules that are covalently linked to RNA were found to be more stable than free NAD (15) at elevated temperatures, suggesting that this process reduces the generation and accumulation of free Nm and ADPR. NAD-RNAs that are converted to ADPR-RNAs via thermal degradation are processed by Saci_NudT5, which releases Nm and 5’-p-RNA.

## Experimental Procedures

### Strains, plasmids, and oligonucleotides

All strains, plasmids, and oligonucleotide sequences used in this study are described in Table S4. This work utilized *S. acidocaldarius* DSM639 MW001, a uracil auxotroph strain (39). Cultures were grown aerobically at 120 rpm and 75°C in Brock medium, pH 3.5 (40). The medium was supplied with 0.1% (w/v) NZ-Amine and 0.2% (w/v) dextrin, and 10 μg/ml uracil. *H. volcanii* was grown as previously described (26). The remaining *E. coli* strains were grown aerobically at 180 rpm and 37°C in LB medium (0.5% (w/v) yeast extract, 1% (w/v) tryptone, 1% (w/v) NaCl). For solid medium, LB medium was mixed with 1.5% (w/v) agar-agar and supplied with the respective antibiotic (0.001% (v/v)). Cell growth was achieved by monitoring the optical density of the cultures at 600 nm.

### RNA extraction and quality control

*S. acidocaldarius* and *H. volcanii* cells were harvested during the mid-logarithmic phase. A 15 ml pellet was lysed with a 2 ml Trizol reagent (Thermofisher), and total RNA was extracted. When needed, total RNA was treated with DNase I (NEB) according to the manufacturer’s instruction and further purified using a Monarch^®^ RNA Cleanup Kit (50 μg) (NEB). RNA integrity was monitored with agarose gels, and RNA concentrations were obtained either with an Implen NanoPhotometer^®^ or with Qubit™ HS RNA assay kit, following the manufacturer’s instructions.

### *In vitro* transcription of NAD-RNAs and ADPR-RNAs

Briefly, each 100 μl IVT reaction contained: a 1 μM DNA template (Supplementary Table 5), 10 μl T7 RNA polymerase (50000 U / ml), 1 mM of each GTP, UTP, CTP, and 4 mM of either NAD or ADPR. The reactions were performed in transcription buffer (40 mM Hepes/KOH pH 8, 22 mM MgCl2, 5 mM DTT) and incubated for 2 hours at 37°C. The transcripts were purified using the Monarch^®^ RNA Cleanup Kit (50 μg) (NEB) and verified on a 6% PAA, 1x TAE, 0.2% APB, and 8M Urea gel.

### LC-MS quantification of NAD

600 μg of total RNA from either *S. acidocaldarius* or *H. volcanii* were divided into six 1.5 ml tubes, 100 μg per tube, and digested with either 10 U of nuclease P1 (NEB) or 10 U of heat-inactivated nuclease P1 for 1 hour at 37°C in a reaction volume of 100 μl. NAD was quantified using a targeted multiple reaction monitoring (MRM) approach in negative ionization mode after chromatographic separation by reversed-phase chromatography according to the following method. The chromatographic separation was performed on an Agilent Infinity II 1260 HPLC system using a YMC C18 column (250 mm, 4.6 mm ID, YMC, Germany) at a constant flow rate of 0.8 ml/min and a constant temperature of 22°C with mobile phase A being 0.4 % acetic acid (Sigma-Aldrich, USA) in water and phase B being 20% Methanol (Honeywell, Morristown, New Jersey, USA) in water. The injection volume was 100 μl. The mobile phase profile consisted of the following steps and linear gradients: 0 – 2 min constant at 0% B; 2 – 16 min from 0 to 35% B; 16 – 13 min from 35 to 100% B; 16 to 20 min constant at 100% B; 20 – 22 min from 100 to 0% B; 22 – 28 min constant at 0% B. An Agilent 6470 mass spectrometer was used in negative mode with an electrospray ionization source and the following conditions: ESI spray voltage 3500 V, sheath gas 400°C at 11 l/min, nebulizer pressure 45 psi and drying gas 170°C at 5 l/min. NAD was identified based on its specific mass transitions (662 (m/z) → 540 (m/z) and 662 (m/z) → 273 (m/z)) and retention time compared to standards. Extracted ion chromatograms of the compound-specific mass transitions were integrated using MassHunter software (Agilent, Santa Clara, CA, USA). Absolute concentrations were calculated based on an external calibration curve.

## Purification of recombinant NUDIX proteins and Saci-aCPSF2

### Saci_NudT5, Saci_NudT5 (NDM), SACI_RS00730, SACI_RS00730 (NDM), SACI_RS00575, and SACI_RS00575 (NDM)

The genes encoding the Nudix proteins SACI_RS00730, Saci_NudT5 and SACI_RS00575 were cloned downstream of the 6x His-tag sequence on the vector pRSFDuet-1 using the restriction sites BamHI and HindIII. NUDIX domain mutant plasmids were generated by performing triple nucleotide exchange via site-directed mutagenesis on the plasmids. The thus generated plasmids were transformed into the expression strain *E. coli* Rosetta 2 DE3 pLysS (Novagen Darmstadt). Cells were grown in a 1l LB medium supplied with 30 μg/ml kanamycin at 37°C, 200 rpm, and protein expression was induced at OD_600nm_= 0.6 – 0.8 with 1 mM IPTG (for SACI_RS00730 and Saci_NudT5) or 0.1 mM IPTG for SACI_RS00575. After further incubation for 3-4 h at 37°C, 200 rpm (for SACI_RS00730 and Saci_NudT5), or overnight at 18°C, 200 rpm for SACI_RS00575, cells were harvested by centrifugation for 15 min at 12.000 x g, 4°C. Pellets were resuspended in 5 ml/g Wash Buffer (WB) (50 mM Tris-HCl, 1 M NaCl, 20 mM Imidazole, 10 mM MgCl_2_, 1 mM DTT, 10% Glycerol, pH 8.0), 1.5 mg lysozyme per gram cells was added to the suspension and incubated on ice for 30 minutes. Next, cells were cracked by sonication, and the supernatant was cleared by centrifugation (20 min at 30.000 x g, RT). Subsequently, the lysate was incubated for 15 min at 75°C, 500 rpm, to denature *E. coli* proteins. After another centrifugation step of 15 min at maximum speed (14.800 rpm, 4°C), the lysate was filtered using a Millex syringe filter (pore size 0.45 μm). One Pierce Centrifuge Column per protein was prepared by washing with several cvs of 20% ethanol and loaded with Ni-NTA Agarose (Qiagen) (stored in 20% ethanol) until each column was filled with ∼2 ml resin. The columns were washed with 10 cvs double-distilled H_2_O (ddH_2_O) followed by 10 cvs Wash Buffer. The lysate was loaded into a resin-filled column, and the flowthrough was saved. Columns were washed with 10 cvs Wash Buffer to remove unspecifically bound proteins. For elution of the His-tagged proteins, the columns were subsequently washed with two times 1 cv of Elution Buffer 1 (WB with 100 mM Imidazole), four times 1 cv of Elution Buffer 2 (WB with 250 mM Imidazole), and three times 1 cv of Elution Buffer 3 (WB with 500 mM Imidazole). Protein elution fractions were analyzed via SDS-PAGE. Protein concentration was analyzed using a Qubit™ 2.0 Fluorometer and the Qubit™ Protein Assay Kit (ThermoFisher Scientific). Finally, proteins were stored at 4°C.

### SACI_RS02625 and SACI_RS02625 (NDM)

*In vitro* protein expression was conducted using the NEBExpress^®^ Cell-free *E. coli* Protein Synthesis System (NEB) according to the manufacturer’s instructions. The plasmids carrying the N-terminally 6x His-tagged genes for SACI_RS02625 and its Nudix domain mutant, which were used as templates for the cell-free expression, were cloned by Genscript Inc. Subsequent purification of the *in vitro* produced proteins was performed using the NEBExpress^®^ Ni Spin Columns (NEB) according to the manufacturer’s protocol.

### Saci-aCPSF2

Recombinant Saci-aCPSF2 was purified under denaturing conditions (8 M Urea) by following standard protocols, using Ni-NTA affinity chromatography (Qiagen) as previously described (32, 31). The purified protein was dialyzed in a storage buffer (100 mM KCl, 50 mM Tris pH 7.0) and stored at -80°C in the presence of 5% glycerol.

### HNudT5

The pET28a-hNudT5 plasmid was transformed into the *E. coli* strain BL21 (DE3). The transformed cells were grown in LB media at 37 °C in the presence of 30 μg/mL kanamycin until OD_600_ reached 0.8. *E. coli* BL21 (DE3) cells were then induced with 0.5 M IPTG, harvested after 3 h, and lysed by sonication (30 s, 50 % power, five times) in HisTrap buffer A (25 mM Tris/HCl pH 8.0, 150 mM NaCl, 5 mM imidazole, 1 mM DTT). The lysate was clarified by centrifugation (14800 rpm, 30 min, 4 °C), and the supernatant was applied to a Ni-NTA HisTrap column (GE Healthcare). The His-tagged protein was eluted with a gradient of HisTrap buffer B (HisTrap buffer A with 500 mM imidazole) and analyzed by SDS-PAGE. Subsequently, enzymes were purified via size exclusion chromatography with a Superose™ 6 Increase 30/100 GL column in gel filtration buffer (25 mM Tris/HCl pH 8.0, 150 mM NaCl). All purified protein samples were 95% pure, judging from SDS–PAGE.

### NAD decapping and ADPR-decapping assays

The NUDIX candidates from *S. acidocaldarius*, NudC (NEB), and HNudT5 were used for decapping assays with NAD- and ADPR-RNAs. Briefly, for each reaction, 1 μl of RNA substrate (15 pmol), 0.5 μl of 10x NEBuffer r3.1, 2.5 μl Nuclease-Free H_2_O and 1 μl of the respective enzyme (15 pmol) were incubated at either 65°C (for the *S. acidocaldarius* NUDIX) or 37°C (for NudC and HNudT5) for 5 minutes. For each sample, a no-enzyme control was established. The reaction was terminated by adding 5 μl of 2x APB-loading buffer (8 M Urea, 10 mM Tris-HCl pH 8, 50 mM EDTA, bromophenol blue, and xylene cyanol blue), and the samples resolved on a 6% PAA, 0.2% APB, 1x TAE, 8 M urea gel. The gel was stained with SYBR™ Gold Nucleic Acid Gel Stain (Thermo Fisher) and visualized with an InstaS GelStick Imager (InstaS Science Imaging™).

### Monitoring of NAD-RNA conversion to ADPR-RNA after heat treatment

Briefly, *in vitro* transcribed NAD-RNA was incubated at 75°C or 85°C for up to 2 hours. Aliquots were taken after 5, 15, 30, 60, and 120 minutes and used as substrates for ADPR-decapping assays with HNudT5, as described above. The reaction was terminated by adding 5 μl of 2x APB-loading buffer, and the samples resolved on a 6% PAA, 0.2% APB, 1x TAE, and 8 M urea gel. The gel was stained with SYBR™ Gold Nucleic Acid Gel Stain (Thermo Fisher) and visualized with an InstaS GelStick Imager (InstaS Science Imaging™). The band intensity was obtained using ImageJ to calculate the half-lives of NAD as previously described (15).

### Saci-aCPSF2 NAD-RNA degradation assay

Degradation assays were carried out as previously described (31). The Saci-aCPSF2 activity was assayed in a 10 μl reaction volume containing 10 mM MgCl_2_, 10 mM KCl, 5 mM Tris pH 7.5, 1.5 pmol of the RNA substrate, and 7.5 pmol (5x excess) of purified Saci-aCPSF2. The reaction mix was incubated from 0 to 90 min at 65°C. The reaction was terminated by adding 10 μl of 2x APB-loading buffer and loaded on 6% PAA, 1x TAE, 0.2% APB, and 8 M urea gel. The gel was stained with SYBR™ Gold Nucleic Acid Gel Stain (Thermo Fisher) and visualized with an InstaS GelStick Imager (InstaS Science Imaging™). The band intensity was obtained using ImageJ (41) to calculate the percentage of remaining RNA after digestion.

### NAD captureSeq library preparation, sequencing, and data analysis

Briefly, as previously described, 600 μg of DNA-free total RNA from each organism was used as input for the preparation of NAD captureSeq libraries (1). Each library was prepared in triplicates (ADPR+ A, B, C, and ADPRC-A, B, and C). Next, PCR products in a range from 150 bp to 300 bp were purified by Bluepippin size selection. The removal of primer dimers was evaluated by using the Agilent DNA 1000 Kit (Agilent) on a Bioanalyzer 2100. The multiplexed library was submitted to NGS on an Illumina HiSeq 3000 or an Illumina MiniSeq in single-end mode and 150 nt read length. Starting Gs of the raw reads and the 3’-adaptor were trimmed using Cutadapt (v2.8), and quality was checked with FASTQC (v0.11.9) (42, 43). Processed reads (≥18 nt) were mapped to the reference genome of either *S. acidocaldarius* or *H. volcanii* using Hisat2 (v2.2.1) (44). After the strand-specific screening, HTSeq was used to count gene hits (45). Statistical and enrichment analyses were performed with DESeq2 (v1.36.0) (22). The Integrative Genomics Viewer (IGV, v2.13.2) was used to inspect and visualize candidate sequences (46). Coding genes were clustered according to their respective arCOGs (47).

### dRNA-seq library preparation, sequencing, and data analysis

To identify transcription start sites (5’-ppp-RNA) in *S. acidocaldarius*, we applied the dRNA-seq technique (48). Briefly, 5 μg of DNA-free total RNA was split into two tubes. One was treated with Terminator™ 5’-Phosphate-Dependent Exonuclease (TEX) (Lucigen, Epicentre) following the manufacturer’s instructions, and the other was submitted to the same reaction but without enzyme. After digestion, the RNA was purified with Monarch® RNA Cleanup Kit (10 μg) (NEB), following the manufacturer’s instructions. Illumina-compatible libraries were prepared using the NEBNext® Small RNA Library Prep Set for Illumina® (NEB). PCR size selection and quality control was performed as described above. The multiplexed library was submitted to NGS using single-end reads, 150 nt read-length on a HiSeq 3000.

For *H. volcanii*, previously published the SRA database (PRJNA324298) and reanalyzed (26). Raw reads were processed as described above. Transcription Start sites were detected using TSSAR: Transcription Start Site Annotation Regime Web Service (v1457945232) (49). Primary transcription start sites matched with NAD-TSS were manually curated using Integrative Genomics Viewer (IGV) (46).

### sRNA-seq and ADPRC+ library comparison

Previously published sRNA-seq (23) data was downloaded from the SRA database (SRX2548838) and reanalyzed. Raw reads were processed as described above. After the strand-specific screening, HTSeq was used to count gene hits (45). Next, genes were ranked according to their fractional representation in each dataset. The top 50 most abundant genes for each library were compared.

### Validation of NGS results with qPCR

Quantitative PCR (qPCR) was performed as described earlier (1) to validate the RNA enrichment observed in the NGS data on the cDNA level. In brief, reactions were performed on a 20 μl scale in duplicate with 3 μl cDNA (1:50 diluted) as a template. qPCR was performed in a Light Cycler 480 instrument (Roche) using the Brilliant III Ultra-Fast SYBRGreen qPCR Mastermix (Agilent). The data were analyzed with the Light Cycler 480 Software (Agilent). Millipore water was used as a negative control, and tRNA-Ile as an internal control gene. The 2-ΔΔCT-method (50) was used to compare APDRC+ sample cDNA with the ADPRC-control cDNA. The primers used for qPCR analysis are listed in Table S4.

### NGS data availability

The generated NGS data are stored at the European Nucleotide Archive (ENA) under the project number PRJEB48624.

## Acknowledgments

We thank Anita Marchfelder for providing *H. volcanii* strains and growth protocols, Sonja-Verena Albers for assistance with *S. acidocaldarius* cultivation and manipulation, Jennifer Kothe for establishing *H. volcanii* growth in our laboratory, Peter Claus for the support during LC/MS experiments, Bruno Huettel for assistance with Illumina sequencing, and Julia Wiegel for technical support. This work was funded by the Deutsche Forschungsgemeinschaft (Heisenberg programme) and the LOEWE Research Cluster “Diffusible Signals”.

## Author Contributions

**J.V.GF:** Experimental design and data analysis. **J.V.GF** and **L.R**.: conceptualization. **R.B:** *Sulfolobus acidocaldarius* Nudix proteins expression and purification. **H.G.MF. A.J**., **A.B**., **and J.S**.: NAD captureSeq library preparation. **N.Po. and K.H**.: HNudT5 purification and assistance with APB-gels. **N.P**.: LC/MS-mediated NAD detection. **J.V.GF**. and **L.R**. wrote the manuscript together with input from all authors.

